# Hyperkalemia, not apoptosis, accurately predicts chilling injury in individual locusts

**DOI:** 10.1101/2020.07.03.186759

**Authors:** Jessica Carrington, Mads Kuhlmann Andersen, Kaylen Brzezinski, Heath MacMillan

**Author notes:** These authors contributed equally to this study (Jessica Carrington, Mads Kuhlmann Andersen).

## Abstract

During prolonged or severe chilling, the majority of insects accrue chilling injuries that are typically quantified by scoring neuromuscular function after rewarming. In the cold, these chill susceptible insects, like the migratory locust (*Locusta migratoria*) suffer a loss of ion and water balance that is hypothesized to initiate cell death. Whether apoptotic or necrotic cell death pathways are responsible for this chilling injury is unclear. Here, we use a caspase-3 specific assay to indirectly quantify apoptosis in three locust tissues (muscle, nerves, and midgut) following prolonged chilling and recovery from an injury-inducing cold exposure. Furthermore, we obtain matching measurements of injury, hemolymph [K^+^], and muscle caspase-3 activity in individual locusts to gain further insight into mechanistic nature of chilling injury. We hypothesized that apoptotic cell death in both muscle and nerve tissue drives motor defects following cold exposure in insects, and that there would be a strong association between cold-induced injury, hyperkalemia, and muscle caspase-3 activity. We found a significant increase in muscle caspase-3 activity, but no such increase was observed in either nervous or gut tissue from the same animals, suggesting that chill injury primarily relates to apoptotic muscle cell death. However, the levels of chilling injury measured at the whole animal level prior to tissue sampling were strongly correlated with the degree of hemolymph hyperkalemia, but not apoptosis. These results support the notion that cold-induced ion balance disruption triggers cell death but also that apoptosis is not the main cell death pathway driving injury in the cold.

**Significance Statement:** Temperature has profound effects on animal fitness and sets limits to animal distribution. To understand and model insect responses to climate, we need to know how temperature sets limits to their survival. There is strong evidence that a collapse of ion and water balance occurs in insects in the cold, and it is generally held that the resulting cold injury is caused by activation of programmed cell death (apoptosis). Here, we directly test this idea and show for the first time that although the loss of ion balance is a strong predictor of individual survival outcomes, apoptosis is not the primary cause of cold-induced injury.

## Introduction

The majority of insects are chill susceptible, meaning they lack physiological mechanisms capable of protecting them from low temperature injury (1). These insects enter a state of paralysis called chill coma (2, 3) that can be reversed following rewarming. The temperature of this paralysis event and the time required to recover the ability to stand following a cold stress (chill coma recovery time; CCRT) are non-lethal and widely used measures of insect chill tolerance (4–7). If a cold exposure is severe enough (defined depending on the species/population under study and its prior thermal history), however, chill susceptible insects suffer from cold-induced injuries - termed chilling injury - that can be sublethal or lethal (1).

Chilling injury typically manifests as defects in an insect’s ability to fly, walk, or stand following chilling, while mortality is often quantified as a complete inability to move, or to undergo a critical phase of development, like adult emergence (3, 8, 9). Thus, although the term chill injury is used to describe multiple organismal outcomes, it most often refers to an insect’s dexterity following cold stress. As such, cell death in the nerves and/or muscles is likely to directly underlie several common cold tolerance metrics.

Cell death is a common consequence of cold exposure in chill susceptible insects, and has been associated with a systemic loss of ion and water homeostasis that occurs during chronic chilling (1). Low temperatures suppress active ion transport (3, 10), and damage paracellular barriers (11–13). During prolonged chilling, a net leak of ions down their concentration gradients across cell membranes and epithelia is commonly observed (8, 14, 15), and a consequence of this mismatch is a systemic rise in extracellular [K^+^] (1, 8, 14–17). The combined effects of slowed active ion transport and elevated extracellular [K^+^] depolarize cells (18–20), triggering excessive calcium influx that is proposed to directly initiate cell death, and both apoptotic and/or necrotic cell death have been blamed for insect chilling injury (18, 21–23).

Understanding when, where, and how cell death occurs in insects during or following chilling is essential to determining the primary causes of organismal chilling injury but is also critical to understanding how insects modulate cold tolerance within the lifetime of an individual (e.g. acclimation) or over evolutionary time. Changes to cold tolerance within an insect appear to arise from physiological adjustments that attenuate the cascade of failure described above (1). For example, cold acclimated individuals and cold-adapted species may rely less on Na^+^ as an extracellular osmolyte (24, 25), better maintain paracellular barrier function in the cold (11, 12), have renal systems more efficient at clearing excess K^+^ from the hemolymph (12, 16, 26, 27), and defend against muscle depolarization induced by low temperatures or elevated hemolymph [K^+^] (20, 28). All of these adjustments serve to protect against injury by targeting upstream causes of physiological failure, but the acquisition of chill tolerance may also be intimately tied to the ability to prevent cell death in the face of homeostatic collapse (23), or even the ability to clear damaged tissue following rewarming (29).

Cellular damage has been repeatedly observed in insect muscles, fat body, and gut epithelia following cold stress, and damage to these organs appears to correlate with chilling injury phenotypes measured at the organismal level (11, 18, 20, 21, 30). These observations of tissue damage, however, have been derived using one of two approaches. First, they have been quantified from live/dead cell viability assays that 1) do not distinguish among necrotic (uncontrolled) and apoptotic (regulated) cell death, and 2) cannot penetrate the blood-brain barrier and thus have not been used to assess nervous damage following chilling (31). With an alternate approach, Yi et al. used a TUNEL assay to quantify DNA fragmentation and interpreted their findings as cell death in the flight muscles of *Drosophila* following chilling occurring primarily via apoptosis (32). Brief pre-exposure to chilling in a manner that improves chill tolerance (a rapid cold-hardening treatment) could inhibit this effect in tissues of flesh flies (*Sarcophaga crassipalpis*) (23). Importantly, however, TUNEL assays cannot distinguish among multiple forms of cell death (33), as DNA fragmentation is a common consequence of cell death. Therefore, apoptosis is likely not acting alone to cause insect chilling injury. Since, the nervous system has not been explored in the context apoptotic or necrotic cell death, whether muscle or nerve damage (or both) cause organismal chilling injury in insect phenotypes remains entirely unclear.

Caspases serve multiple functions in insects (34, 35), but their primary role is in programmed cell death cascades where they are produced in advance of cell death and maintained in an inactive precursor form (pro-caspase). Regulated cell death pathways are generally well-conserved among animals, and the roles of individual caspases are increasingly well-understood (36). In *Drosophila*, Drice (a caspase-3 ortholog) is the major executioner caspase that is essential for programmed cell death during development and in response to tissue/cell damage (37–41). This central role of caspase-3 and its orthologs as important effectors driving cell destruction is conserved among many species, including insects and mammals. Because caspase- 3 and its orthologs appear to be mainly associated with apoptotic cell death and not necrosis (42), it can be a useful tool for understanding the ultimate causes of chilling injury.

Here, we use the migratory locust (*Locusta migratoria*) to test the hypothesis that ionoregulatory collapse drives caspase-mediated cell death in both the nerves and muscles and is responsible for insect chilling injury. We exposed locusts to up to 48 h at -2°C to determine a duration of exposure that caused significant and variable sub-lethal chilling injury and used this treatment to examine activation of caspase-3-like proteins (executioner caspases associated with apoptosis) in a thoracic muscle, the metathoracic ganglion, and the midgut (as a negative control as midgut cells use autophagy, not caspase activation for programmed cell death (43)). Since caspase-3 activation occurred specifically in the muscles in the cold, we obtained matching measurements of survival, hemolymph [K^+^], and muscle executioner caspase activity from individual locusts during cold exposure. This allowed us to investigate the links between these parameters and generate the first data relating individual variation among these measures in any insect. With this approach we provide evidence that injury to the muscles, and not the nerves, is most likely responsible for motor defects following cold exposure, and that while cold stress activates muscle caspase, the degree of hyperkalemia is a far better quantitative predictor for organismal chilling injury than muscle executioner caspase activity. Thus, other cell death pathways are likely responsible for chilling injury.

## Results

### Chill coma recovery time and survival following exposure to -2°C

The cold tolerance of locusts was examined by measuring chill coma recovery time (CCRT) at specific time points during exposure to -2°C and was followed by a survival assessment (scale of 0-5) 24 h after the end of the cold exposure (Fig. 1). Exposure to -2°C gradually increased CCRT for both sexes (t_2,21_ = 13.8, P < 0.001 for exposure time; t_1,21_ = 0.8, P = 0.446 for sex), however, females became increasingly slower at recovering as exposure time increased (interaction: t_2,21_ = -2.8, P = 0.010) such that recovery took 9.2 ± 0.3 min and 9.0 ± 0.4 min for females and males, respectively, after 2 h of exposure and increased to 49.9 ± 3.9 min and 36.6 ± 9.2 min after 24 h. After 48 h no locusts recovered within the 60 min time limit (Fig. 1A). A similar decrease in post-exposure performance was found for the survival scores (no effect of sex); survival scores decreased from 4.9 ± 0.1 after 2 h of cold exposure to 1.0 ± 0.3 after 48 h (H = 25.5, P < 0.001; Fig. 1B).

**Figure 1.**
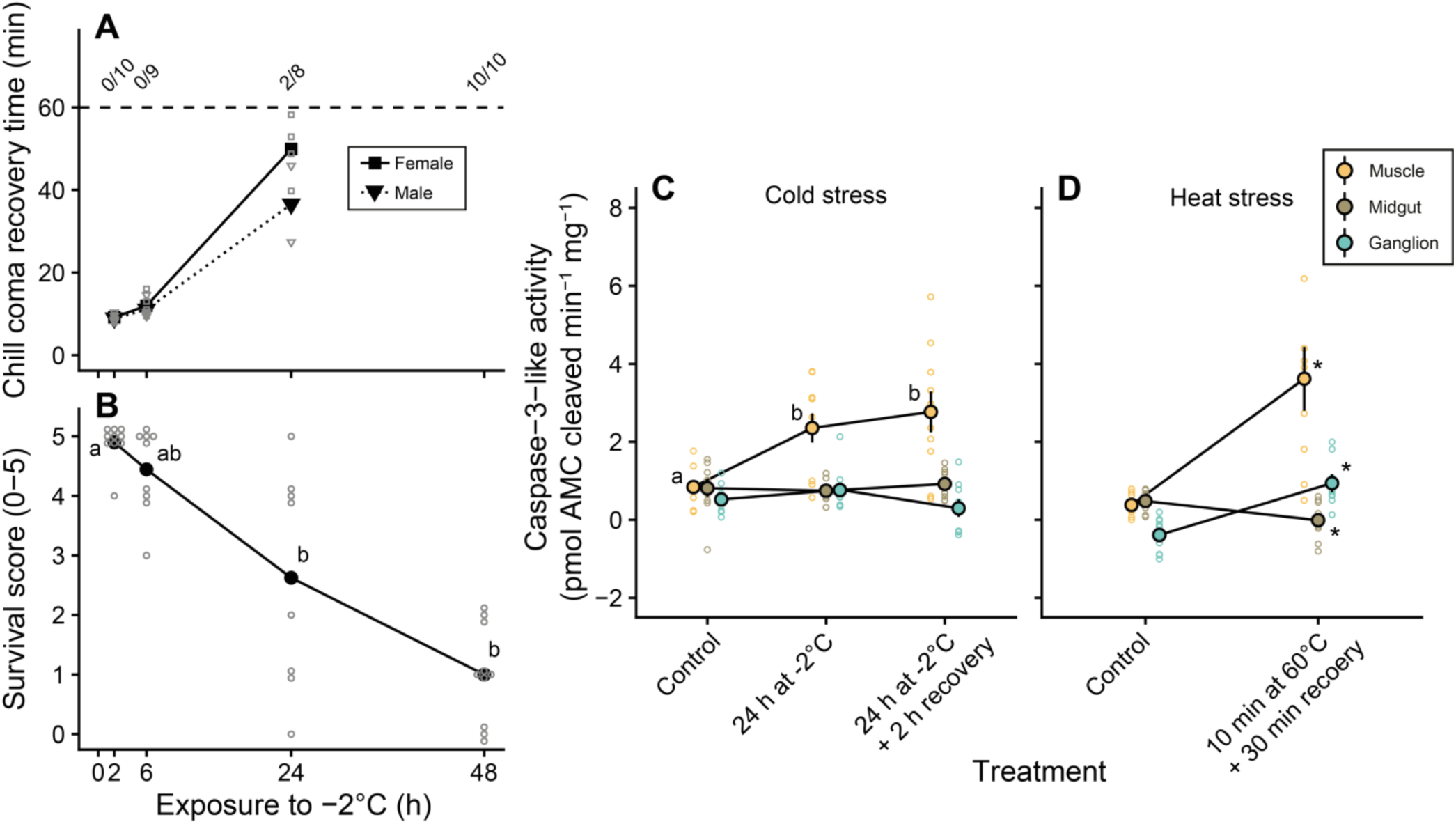
Cold stress that causes injury also causes activation of caspase-3-like activity in the muscles of locusts. Prolonged exposure to -2°C gradually (A) increased the time needed for locusts to assume a standing position in both females (squares) and males (triangles) and (B) reduced the survival outcome after recovery at a permissive temperature. (C) During this cold exposure caspase-3-like was increased in muscle tissue (orange), but remained the same in midgut (brown) and nervous tissue (blue). (D) Lethal heat exposure was used as a positive control, and resulted in caspase-3-like activation in muscle and nervous tissue, while caspase-3- like activity decreased in the midgut. Individual data points are represented by small, empty symbols. Error bars not visible (for C and D) are occluded by the symbols.

### Caspase-3 activity induced by exposure to thermal extremes

To test whether the observed reduction in survival was related to an increase in apoptotic activity, we measured caspase-3-like activity in muscle (flight muscle M90, after Snodgrass (44)), nervous tissue (metathoracic ganglion), and midgut (negative control) both after an intermediate cold exposure and after a brief recovery period, and followed each up with a positive heat exposure control (see Fig. 1). Exposure to -2°C for 24 h increased caspase-3-like activity in muscle tissue from 0.8 ± 0.2 pmol AMC cleaved min^-1^ mg^-1^ in control locusts to 2.4 ± 0.4 pmol AMC cleaved min^-1^ mg^-1^, which was similar to the 2.8 ± 0.5 pmol AMC cleaved min^-1^ mg^-1^ measured after 2 h of recovery (F_2,26_ = 6.4, P = 0.006; Fig. 1C). Caspase-3-like activity remained unchanged in both midgut tissue and nervous tissue (F_2,24_ = 0.3, P = 0.755 and F_2,23_ = 1.6, P = 0.228, respectively) with activities ranging from ∼ 0.3 t 0.9 pmol AMC cleaved min^-1^ mg^-1^ (Fig. 1C). Brief exposure to 60°C was used a positive control for caspase activation (Fig. 1D), and increased caspase-3-like activity in flight muscle from 0.4 ± 0.1 pmol AMC cleaved min^-1^ mg^-1^ to 3.6 ± 0.8 pmol AMC cleaved min^-1^ mg^-1^ (t_16_ = -3.9, P = 0.001). Unlike the cold, lethal heat stress also increased caspase-3-like activity in nervous tissue from -0.4 ± 0.2 pmol AMC cleaved min^-1^ mg^-1^ to 0.9 ± 0.2 pmol AMC cleaved min^-1^ mg^-1^ (t15 = -4.9, P < 0.001), while it decreased in midgut tissue from 0.5 ± 0.1 pmol AMC cleaved min^-1^ mg^-1^ to 0.0 ± 0.2 pmol AMC cleaved min^-1^ mg^-1^ (t_14_ = 0.036).

### Individual variation in survival, hemolymph K^+^ concentration, and caspase-3 activity

To gain further insight into the relationship between survival, ion balance, and caspase-3-like activity, we took advantage of the wide inter-individual variation noted in these variables in the first set of experiments. Here, we scored survival and measured hemolymph K^+^ concentration and flight muscle caspase-3-like activity in the same individuals, using unexposed locusts and locusts exposed to 24 and 48 h of exposure to -2°C (and 2 h of recovery, Fig. 2). As previously demonstrated, survival score decreased with longer cold exposures (H = 36.6, P < 0.001, Fig. 2A). In the same locusts, hemolymph K^+^ concentration increased during exposure and recovered over the two hours of recovery before dissection of the muscle tissue (F_5,97_ = 51.1, P < 0.001, Fig. 2B). Specifically, hemolymph [K^+^] increased from 9.7 ± 0.6 mmol L^-1^ in controls to 23.8 ± 0.8 mmol L^-1^ after 24 h and was restored to 16.0 ± 0.8 mmol L^-1^ after recovery. In the group exposed for 48 h, it increased to 37.0 ± 1.4 mmol L^-1^ and returned to 24.6 ± 1.6 mmol L^-1^ after the recovery period. Correlating survival score and hemolymph [K^+^] for each locust revealed a tight, sigmoidal-like relationship with an IC_50_ (“Injury Concentration 50”; hemolymph [K^+^] that correlates to a 50% reduction in survival score) of 34.8 ± 0.8 mmol L^-1^ (Fig. 2C). In the same animals, muscle caspase-3-like activity increased during cold exposure (samples taken after the 2 h recovery period) from -0.9 ± 0.2 pmol AMC cleaved min^-1^ mg^-1^ to 7.3 ± 2.2 and 5.3 ± 2.1 pmol AMC cleaved min^-1^ mg^-1^ after 24 and 48 h, respectively (H =16.8, P < 0.001, Fig. 2D). Correlating survival scores and muscle caspase-3-like activities revealed no relationship between these parameters (linear regression: t_1,48_ = -0.7, P = 0.473; see Fig. 2E). One would expect that flight muscle caspase-3 activity would correlate better with the wing-specific score, and although the correlation was stronger, the relationship did not reach statistical significance (t_1,48_ = -1.7, P = 0.089, see Fig. S1). Furthermore, there was no relationship between caspase-3-like activity and hemolymph K^+^ concentration (linear regression: t_1,48_ = 1.0, P = 0.326, correlation not shown). The poor predictive power of muscle caspase-3-like activity is likely partially caused by the large variation in activity; a minority of muscle samples from cold exposed locusts have very high caspase-3-like activity (>10 pmol AMC cleaved min^-1^ mg^-1^). When these samples are removed (using Grubb’s test for outliers), all correlations became statistically significant using linear regression (apoptosis vs. survival score: t_1,39_ = -2,9, P = 0.005, R^5^ = 0.164; apoptosis vs. wing score: t_1,39_ = -4.4, P < 0.001, R^2^ = 0.316, apoptosis vs. hemolymph [K^+^]: t_1,39_ = 3.1, P = 0.003, R^2^ = 0.178; see Fig. S2). Taking this approach, however, 1) reduces our sample size to a degree we find uncomfortable (nine outliers out of 50 data points removed), and 2) yields relationships between muscle caspase activity and survival scores that, while significant, still do not come close to reaching the explan<u>a</u>tory power of hemolymph [K^+^]. We therefore opted to retain the entire dataset in Fig. 2.

**Figure 2.**
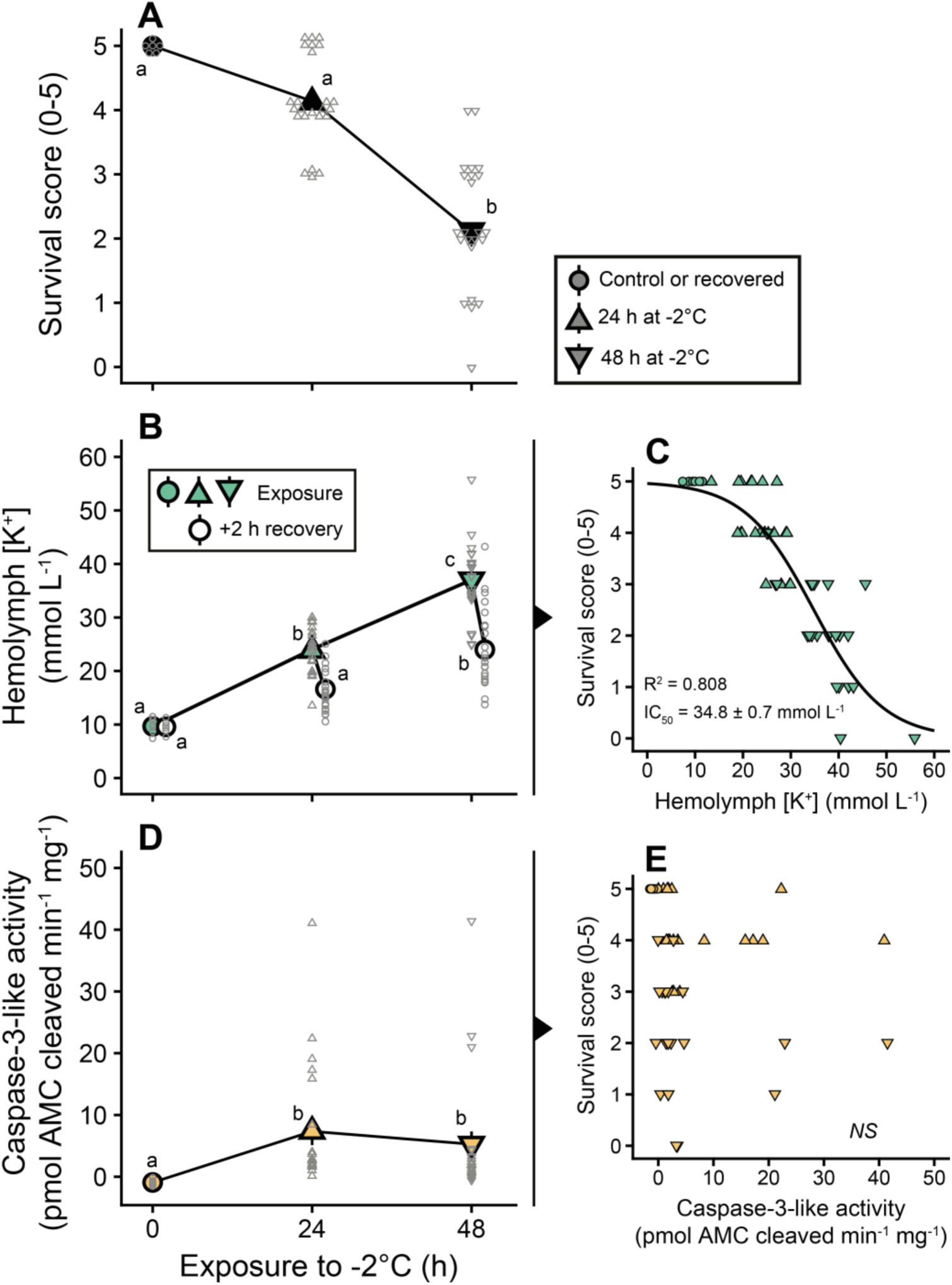
Individual variation in cold-induced hyperkalemia predicts individual survival outcomes while caspase-3-like activity in the muscles does not. (A) Exposure to stressful cold reduces survival and (B) increases hemolymph [K^+^] (hyperkalemia) with (C) a strong sigmoidal correlation between the two 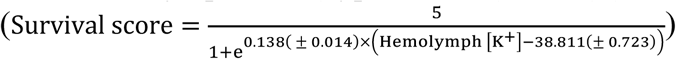. (D) Caspase-3-like-mediated apoptosis was activated during the same exposure, but (E) did not correlate with the survival score.

## Discussion

### Stressful cold causes injury and activates programmed cell death in muscle tissue

Like other chill susceptible insects, locusts sustain injuries during cold exposure (1), but the physiological mechanisms underlying the onset of these chill–related injuries remain elusive. We designed the present study to investigate whether chilling injuries could be caused by cold-induced activation of a common cell death pathway, namely caspase-3-mediated apoptosis. Furthermore, we tested whether the degree of chill injury was correlated with levels of caspase-3 activity and/or ion balance disruption in individual locusts.

As has been previously demonstrated, prolonged exposure to -2°C causes injury in locusts in a time-dependent manner, both in terms of a slowed recovery time and less favourable survival outcome after exposure (Fig. 1A,B; (18–21)). Our way of quantifying chill injury in the present study is based on the ability of locusts to perform coordinated movements after cold exposure (i.e. the ability to move immediately after coma or after a recovery period), and the behavioural deficits after exposure could therefore stem from 1) debilitating injury to the muscles themselves, 2) injury to the integrating neural centers, 3) loss of function in the neuromuscular excitation-contraction coupling (not investigated here), or 4) a combination of all three (1, 45).

Cold-induced cell death in insect muscle is thought to be the consequence of a debilitating cascade, at the centre of which is a loss of ionoregulatory capacity that drives hemolymph hyperkalemia. This hyperkalemia, in turn, depolarizes muscle tissue and induces an excessive Ca^2+^ influx, increasing the intracellular [Ca^2+^], and this is thought to activate apoptotic/necrotic pathways and thereby drive injury phenotypes (1, 18, 21–23, 45). In our experiments we found that exposure to both prolonged cold and lethal heat (positive control) induced a marked increase in caspase-3-like activity in muscle tissue (Fig. 1C,D and Fig. 2D). Caspase-3 is one of the main executioner caspases responsible for programmed cell death, and while effector caspases can be activated by several up-stream initiator caspases, caspase-3 in particular appears to be mainly associated with apoptotic rather than necrotic cell death (42), thus we demonstrate that muscle cell death caused by stressful temperatures is at least partially caused by caspase-3-mediated apoptosis. This is supported by the findings of Yi and Lee who demonstrated that cold-induced cell death in *D. melanogaster* was associated with DNA fragmentation (32), a common marker for cell death. The exposure used to induce cell death in the present study causes hemolymph hyperkalemia and muscle membrane depolarization (Fig. 2D; MacMillan et al., 2014), thus our findings support a link between cold-induced ionoregulatory collapse and cell death (21, 22). Interestingly, the locust gut is also injured by hemolymph hyperkalemia (21), however, we found no increase in caspase-3-like activity in the midgut in response to cold exposure (Fig. 1C). We noted a small but statistically significant decrease (rather than the expected increase) in caspase- 3-like activity in the midgut after severe heat exposure. What, if anything, drove this small effect is unclear. Together, these results from our cold and heat-stress experiments suggest that unlike muscles and nervous tissue, cell death does not occur through activation of caspase-3 orthologs in the midgut of locusts, which is similar to what has been established for *Drosophila* (43).

### A lack of cold-induced apoptotic cell death in the central nervous system

Loss of coordinated movements after cold exposure can, as mentioned above, be caused by cold-induced injury to the integrating centres in the nervous system. To estimate injury to the central nervous system (CNS) we measured caspase-3-like activity in the metathoracic ganglion, and found increased activity only after exposure to lethal heat (Fig. 2C,D). This differs from the muscle tissue where both heat and cold initiated caspase-3-mediated cell death. One possible explanation for this lies in the differential distribution and abundance of Ca^2+^ channels in insect nerve and muscle tissue: Insect muscles use Ca^2+^ ions for action potential generation and have a high and relatively even distribution of voltage-gated Ca^2+^ channels resulting in the high Ca^2+^ currents necessary muscle excitation, whereas insect nerves use Na^+^ channels for action potential generation and have highly localized Ca^2+^ channel distribution resulting in lower whole-cell currents (46–48). Thus, if the onset of chilling injury is based purely on depolarization-mediated Ca^2+^ entry, tissue injury could in principle be driven entirely by the presence or absence of voltage-gated Ca^2+^ channels. This is supported by the finding that blockade of Ca^2+^ channels can prevent the onset of chilling injury (21).

The central nervous system not only distinguishes itself from muscle on the basis of Ca^2+^ channel distribution, but also differs in its physiological response to stressful conditions: During exposure to thermal extremes the CNS undergoes a phenomenon known as a spreading depolarization (SD) (49, 50). SD events are characterized by a rapid surge in interstitial [K^+^] that completely silences the CNS at a temperature closely associated with the loss of coordinated movements at the CT_min_ and CT_max_ (4, 5, 51). However, while an increase in extracellular [K^+^] in the hemolymph appears to be detrimental, the SD event has been hypothesized to serve a neuroprotective function in insects (7, 51). Indeed, it has been proposed that the large shifts in interstitial ion concentrations that occur during SD (not only [K^+^] changes, see (52)) could induce channel and/or spike arrest in the CNS such that the SD serves to lower metabolic demand during exposure to extreme conditions (53–55). Furthermore, it was recently suggested that SD events themselves are benign unless occurring in metabolically compromised tissues (56). Exposure to extreme heat severely challenges aerobic metabolism in insects while energy balance is generally maintained during cold exposure (57, 58), and our finding that heat, and not cold, increases caspase-3-mediated cell death in the locust CNS therefore at least partially supports an adaptive nature of SD events.

The hypothesis that cold-induced SD is protective in insects is indeed appealing and has some degree of support from our data, as only muscle appeared to suffer apoptotic cell death during the cold exposure. However, it is also possible that the CNS suffers injury *via* other pathways. Specifically, Boutilier (22) proposed that cold-induced cell death could occur via cell swelling-induced necrosis (see (59)) in rat glial cells and it is therefore possible that the CNS (in the ganglia or elsewhere) suffers considerable injury that simply cannot be detected with a caspase-3 assay.

### Individual variation in hemolymph [K^+^] predicts survival outcomes during cold exposure

The capacity to prevent the systemic loss of ion and water homeostasis during cold exposure is thought to underlie the ability to tolerate prolonged cold exposures and avoid injury (1, 45). Until now, however, no study has quantified the degree of chilling injury and ion balance disruption in the same individual of any insect species. We took advantage of the variation in survival outcome in cold-exposed locusts to investigate the role of individual variation in ionoregulatory capacity in facilitating cold tolerance by measuring survival outcome, hemolymph [K^+^], and caspase-3-like activity in the muscles of individual locusts (Fig. 2). As before, we found that poor survival outcomes were generally associated with hemolymph hyperkalemia, but we also found a strong, negative sigmoidal relationship between the degree of chilling injury and degree of hyperkalemia (Fig. 2A-C). Thus, our findings provide strong support for a link between ionoregulatory capacity and cold tolerance on the level of individual insects. In the current model for insect chilling injury, cell death is initiated by a cold- and hyperkalemia-mediated depolarization of muscle membranes that *via* catastrophic Ca^2+^ overload activates apoptotic and/or necrotic pathways (1, 21, 45), so we expected that muscle capase-3-like activation would be similarly correlated with hyperkalemia and survival outcomes. Surprisingly, however, in spite of finding that caspase-3-like activity was increased in cold exposed (and hyperkalemic) locusts (Fig. 2D), we found no relationship between caspase-3-like activity and survival score (Fig. 2E). The same was true for caspase-3-like activity and wing score, and caspase-3-like activity and hemolymph [K^+^] (Fig. S1).

The current model for chilling injury implicates Ca^2+^ as a key signalling molecule in activating apoptosis (21, 30), however, increased cytosolic [Ca^2+^] also activates other cell death pathways such as autophagy and necrosis (59–61). It is therefore likely that not all cell death in locust muscle is driven by caspase-3-like activity, or even by apoptosis. As mentioned earlier, Boutilier (22) proposed that cell swelling could contribute to cold-induced cell death and it has been shown by Denton et al. (43) that cell death in the midgut of *Drosophila melanogaster* mutants was caused primarily by autophagy. Thus, it is likely that other cell death pathways play more critical roles in the cold-induced cell death that has observed in insect muscle using live/dead assays (18, 20, 21). Indeed, damage to the cell membrane (utilized by live/dead assays to estimate viability) is a phenomenon commonly associated with necrosis caused by cell swelling (59). It therefore seems likely that the tight link between hemolymph hyperkalemia and cell death (18, 21) is based on, or at least includes, observations of necrotic cell death.

Our inability to correlate caspase-3-like activity with survival outcomes could alternatively be explained by the use of a single flight muscle as a sample to predict injury at the organismal level. Some support for this can be found in the slightly stronger (but still not statistically significant) association between the wing-specific survival score and muscle caspase-3-like activity (see Fig. S1). Lastly, the possibility remains that the mechanism underlying cold-induced behavioural deficits is not associated with cell death, but *via* other detrimental effects of cold and/or hyperkalemia on the neuromuscular systems, for example, cold exposure has been shown to affect synaptic function in *Drosophila melanogaster* and the crayfish *Procambarus clarkia* (62), and disruption of synaptic function in cold stressed animals could similarly serve to explain neuromuscular injury following rewarming.

## Conclusions

Overall, our findings suggest that cold stress activates apoptotic signaling cascades in the muscles, but not nervous tissues of a chill susceptible insect. Hyperkalemia has been repeatedly observed as a consequence of chilling in insects, and we found for the first time that it is a strong predictor of individual neuromuscular defects following rewarming. Although cold activates apoptosis in the muscles of locusts, caspase activity does not correlate with individual organismal injury phenotypes. We argue that hemolymph K^+^ is a better predictor of chilling injury primarily because 1) K^+^ imbalance is central to determining whether or not an insect is injured and 2) other cell death pathways (most likely necrosis) are at play. To integrate these new findings into our current understanding of chilling injury we present a revised model of the mechanisms driving organismal chilling injury in chill susceptible insects (Fig. 3), which highlights the critical importance of distinguishing among apoptosis and other forms of cell death in furthering our understanding of insect cold tolerance. Only by doing so can we understand how cold adapted species and populations can avoid and repair cellular damage during and following cold stress.

**Figure 3.**
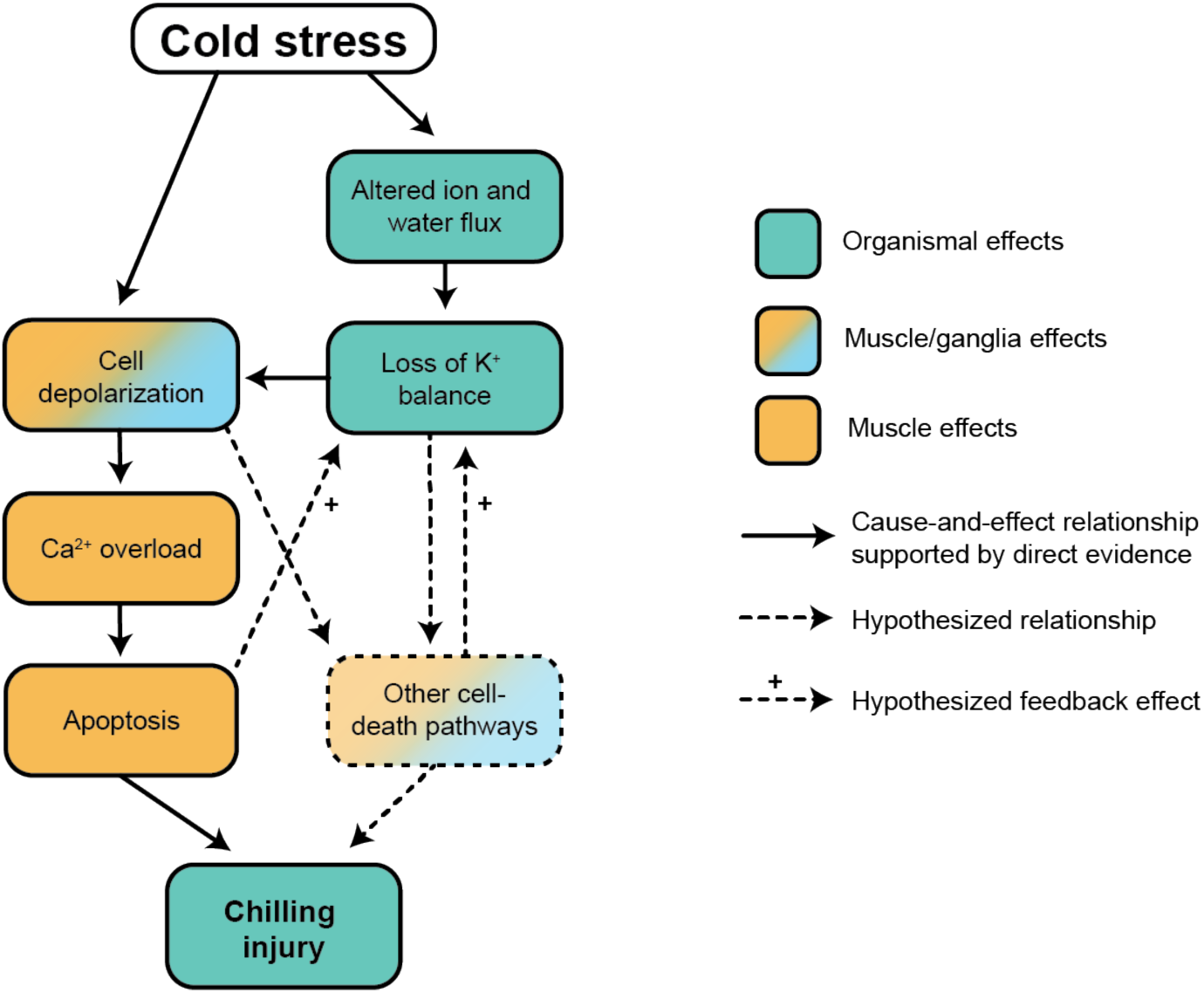
A revised model of cause-and-effect relationships between cold exposure and chilling injury phenotypes in insects. Exposure to stressful cold directly depolarizes cell membranes, and this effect is exacerbated by both a systemic (hemolymph; impacting muscles) and local (spreading depolarization; impacting the central nervous system) loss of K^+^ balance. This causes cell membrane depolarization that drives a catastrophic increase in cytosolic [Ca^+^] in muscle cells which activates executioner caspases and subsequent apoptotic cell death leading to some injury at the organismal level. Based on the findings of the present study, however, it is likely that other cell-death pathways (e.g. necrosis) or deleterious (and likely Ca^2+^-overload-independent) mechanisms are activated by membrane depolarization and cause further chilling injury.

## Materials and Methods

### Animal husbandry

Our colony of *Locusta migratoria* is maintained at Carleton University, Ottawa, ON. This colony is continuously breeding under crowded conditions. Locusts are held at 30°C, with a 16:8 day/night cycle, fed on wheatgrass and an oat mixture (65% oats, 10% wheat germ, 10% wheat bran, 5% skim milk powder). For all experiments, locusts were taken from a crowded cage at 3-4 weeks post-final ecdysis, and were used in a ∼ 1:1 sex ratio for all experiments.

### Chill coma recovery time and survival following exposure to -2°C

Chill coma recovery time and chilling injury were assessed following exposure to -2°C following previously described methods (13). Each locust was placed in a 50 mL ventilated polypropylene centrifuge tube before being placed in a mixture of ethylene glycol and water (with holes in the tube lid in contact with the air) inside a refrigerated circulator (28L with advanced programmable controller, VWR International, Radnor, USA). Temperature was set to hold locusts at 20°C for 15 min and then decrease to -2°C at a rate of -0.1°C min^-1^ and held there for up to 48 h. Groups (N = 10 per group) of locusts were removed from the bath at four time points (2, 6, 24, and 48 h), and a control group was held in tubes at room temperature (∼22°C) for 24 h. The control group was not fed nor allowed to drink for the entire 24 h to best match the experimental groups. Cooling bath temperature was confirmed to keep locusts at -2°C (± 0.5°C) using three type-K thermocouples (connected to a TC-08 data logger, Pico Technology Inc., St. Neots, Cambridgeshire, UK) in three different tubes containing locusts.

Once removed from the cooling bath, locusts were placed at room temperature (22 ± 0.5°C) and gently stimulated every five minutes until they were observed to stand, or until 60 min had passed. Locusts were then returned to their respective 50 mL tube, with access to food and water, until survival score was assessed 24 h later. Survival score was rated on a scale of 0-5 in a manner similar to that used previously (19) by removing each locust from the tube and gently coaxing them to move. Survival was scored as follows: 0 = motionless/dead, 1 = twitching without coordinated movement, 2 = able to move but unable to stand, 3 = able to stand, 4 = able to walk, jump, and initiate flight, but with slow reaction time, 5 = able to walk, jump and initiate flight with no observable defects or delays in reaction time.

### Caspase-3 activity following cold exposure

Caspase-3-like activity was measured in three tissues dissected from locusts from three treatment groups (N = 6 per treatment): 1) Controls held at 28°C for 24 h, 2) cold exposed and dissected immediately after 24 h at -2°C, and 3) cold exposed and dissected after a 2 h recovery period to test for delayed activation of caspase-3. The cooling bath followed an identical ramping regime used to assess chill coma recovery and chilling injury.

To isolate tissues, locusts were quickly decapitated, and all appendages were removed before a single incision was made in the anterior-posterior axis of the dorsal cuticle. The body cavity was pinned open, submerged in standard locust saline (in mmol L^-1^: 140 NaCl, 8 KCl, 3 CaCl_2_, 2 MgCl_2_, 90 sucrose, 5 glucose, 5 trehalose, 1 proline, 10 HEPES; pH 7.2), and a sample of the posterior midgut (excluding the caeca) was taken and cleaned with an aliquot of clean saline. Then the posterior metathoracic tergocoxal muscle (M90 following (44), a flight muscle) and the metathoracic ganglion were dissected out. All tissues were snap frozen in liquid nitrogen after dissection and stored at -80°C until use.

Caspase-3-like activity was quantified using the EnzChek Caspase-3 Assay kit #1 (Molecular Probes, Eugene, OR, USA). Tissue samples were thawed on ice for 5 min, before being suspended in 100 μL/mg lysis buffer (10 mmol L^-1^ TRIS; pH 7.5, 0.1 mmol L^-1^ NaCl, 1 mmol L^-^ 1 EDTA, 0.01% Triton X-100, in dH_2_O). Each sample was sonicated for rounds of 5 s (with 15 s breaks on ice between rounds to prevent overheating) until fully homogenized. Samples were then centrifuged for 5 min at 2000 × *g* at 5°C. A 50 μL aliquot of sample supernatant was transferred to a black, clear bottomed, 96-well microplate.

Along with blank samples (containing only 100 μL lysis buffer), two additional controls were run in each assay plate. First, a subset of samples containing 1 μL of (1 mmol L^-1^ in DMSO) Ac-DEVD-CHO (a specific inhibitor of caspase-3-like proteases) were included in a subset of duplicate wells to confirm that the fluorescence observed was specifically caused by the activity of caspase-3-like proteases (confirmed). Secondly, samples with 1 μL of the DMSO solution were measured to control for the effect of the DMSO itself (there was none).

A 2x working solution was prepared by adding 2% V:V Z-DEVD-AMC substrate (10 mmol L^-1^ in DMSO) to the 2x reaction buffer (2.5 mmol L^-1^ PIPES, 0.5 mmol L^-1^ EDTA, 0.025% CHAPS, diluted in dH2O, pH 7.4, and 1% V:V DTT (in 1 mmol L^-1^ in DMSO)). 50 μL of the working solution was added to each sample and control (combined volume of 100 μL). The samples and controls were left to incubate for 30 min at room temperature. To quantify caspase-3 activity through the DEVD-AMC substrate, serial dilutions of AMC ranging from 0-100µM (from a stock solution also containing 10 mmol L^-1^ DMSO) were added to single wells (100 µL each). Fluorescence of the samples (excitation: 324 nm, emission: 441 nm) was measured with a CYTATION5 fluorescence spectrophotometer (BioTek Instruments, Winooski, VT, USA).

### Heat shock controls

We were surprised to observe differences in caspase-3-like activity between the nerve and muscle tissues following chilling, so we examined whether this was a general pattern following thermal stress that causes organismal injury or was specific to our chilling protocol. We thus purposefully induced apoptosis in a separate group of locusts (N = 9) by exposing them to a lethal heat shock (60 ± 1°C for ∼ 10 min). After resting at 28°C for 30 minutes, the locusts were dissected. While not all locusts were completely motionless directly after the heat shock, all of the locusts were scored as a 0 (dead/motionless) after the 30 min recovery period. The dissection and caspase detection protocol described above was then repeated for all three tissues collected from these locusts.

### Matching measurements of injury, hemolymph K^+^ concentration, and muscle caspase-3 activity

In a separate set of experiments, locusts were exposed to -2°C for 0, 24, and 48 h (following the same procedure as above; the 0 h group was never exposed) after which they were moved to room temperature. Immediately after removal from the cold, a small hemolymph sample was taken by gently penetrating the neck membrane between the head and the thorax with a glass capillary tube and having the tube collect approximately 1 μL of hemolymph. The hemolymph was then transferred to a small dish and kept under hydrated mineral oil. After 2 h of recovery, the locusts were scored for survival (0-5 as described above) and an additional wing-specific score was estimated (also 0-5) to rank motor function defects and injury to the wing muscles (0 = appendage motionless, 1 = twitching, 2 = slightly reactive, 3 = reaction to stimulus, limited range of motion, 4 = full range of motion, but uncoordinated, or with delayed reaction, 5 = fully functional). After scoring locusts, a second hemolymph sample was taken, and the M90 flight muscle was dissected out under standard saline, quickly blotted dry, transferred to a pre-weighed Eppendorf tube and weighed, snap-frozen in liquid N2 and stored at -80°C until measurement of caspase-3-like activity (as described above).

Hemolymph [K^+^] was measured using ion-selective glass microelectrodes as described by (16). Briefly, glass capillaries (TW-150-4, World Precision Instruments (WPI), Sarasota, FL, USA) were pulled to a fine tip and silanized in an atmosphere of N,N-dimethyltrimethylsilylamine (Sigma Aldrich, St. Louis, MO, USA). Silanized glass microelectrodes were then back-filled with 100 mmol L^-1^ KCl and front-filled with K^+^ ionophore (K^+^ ionophore I, cocktail B, Sigma Aldrich, St. Louis, MO, USA). A thinly pulled glass electrode (IB200F-4, WPI) back-billed with 500 mmol L^-1^ KCl was used as a reference. Before every measurement, electrodes were calibrated in 10 and 100 mmol L^-1^ KCl solutions (LiCl was used to balance osmolality) to obtain the Nernstian slope (∼ 58.2 mV per 10-fold change in concentration at 25°C), and only electrodes with a slope between 50 and 62 mV were used (mean ± standard deviation of 21 electrodes: 54.3 ± 2.4 mV). For this experiment 6 locusts were used as controls and 24 locusts were exposed for both the 24 h and 48 h. It was not possible to obtain a second hemolymph sample from five locusts (three and two from the 24 h and 48 h exposure group, respectively, so the sample size here was N = 6, 21, and 22), and four muscle samples were lost during transfer out of the liquid N2 (three and one from the 24 h and 48 h exposure group, respectively, lowering the sample size for muscle caspase-3-like activity to N = 6, 21, and 23).

### Data analysis

All data analysis was completed in R version 3.5.3 (63). All datasets were tested for normality using boxplots and Shapiro-Wilk tests (shapiro.test() function), and non-parametric approaches were used when appropriate. All starting models included sex as a factor, but this factor was eliminated in all but one case where it interacted with exposure time: Chill coma recovery times following exposure to -2°C were analysed using a generalized linear model with exposure time as a continuous variable and sex as a factor. Survival scores were compared among exposure times using Kruskall-Wallis tests followed by Dunn’s multiple comparisons tests using the kruskal.test() and dunnTest() (FSA package) functions, respectively. The effect of cold exposure on caspase-3 activity was analysed using separate one-way ANOVAs for each tissue, followed by Tukey HSD *post hoc* tests. Heat-activated caspase-3 activity (i.e. the positive control) in each tissue was compared to controls using t-tests. For the dataset on individual variation, the effect of cold exposure on the survival score and caspase-3 activity were analysed using Kruskal-Wallis tests followed by Dunn’s multiple comparison tests, while those of hemolymph K+ concentration was analysed using a one-way ANOVA followed by Tukey’s HSD *post hoc* test. Correlations between survival scores and caspase-3 or hemolymph K+ concentration were tested using linear regression and non-linear regression to a sigmoidal model (using the nls() function; model parameters specified in figure text), respectively, and the best fitting model (based on R2 values and AIC scores), if statistically significant, is displayed. All values listed are means ± s.e.m. unless otherwise stated, and the critical level for statistical significance was 0.05 in all analyses.

## Supporting information

Data Archive

## Acknowledgements

The authors wish to thank Marshall Ritchie for taking care of the locust colony during the time this research was being conducted.

## Data Availability

All data is provided as a supplementary file for review and the same file will be included as supplementary material should the manuscript be accepted for publication.

