## Supplementary figures for "Hyperkalemia, not apoptosis, accurately predicts chilling injury in individual locusts"

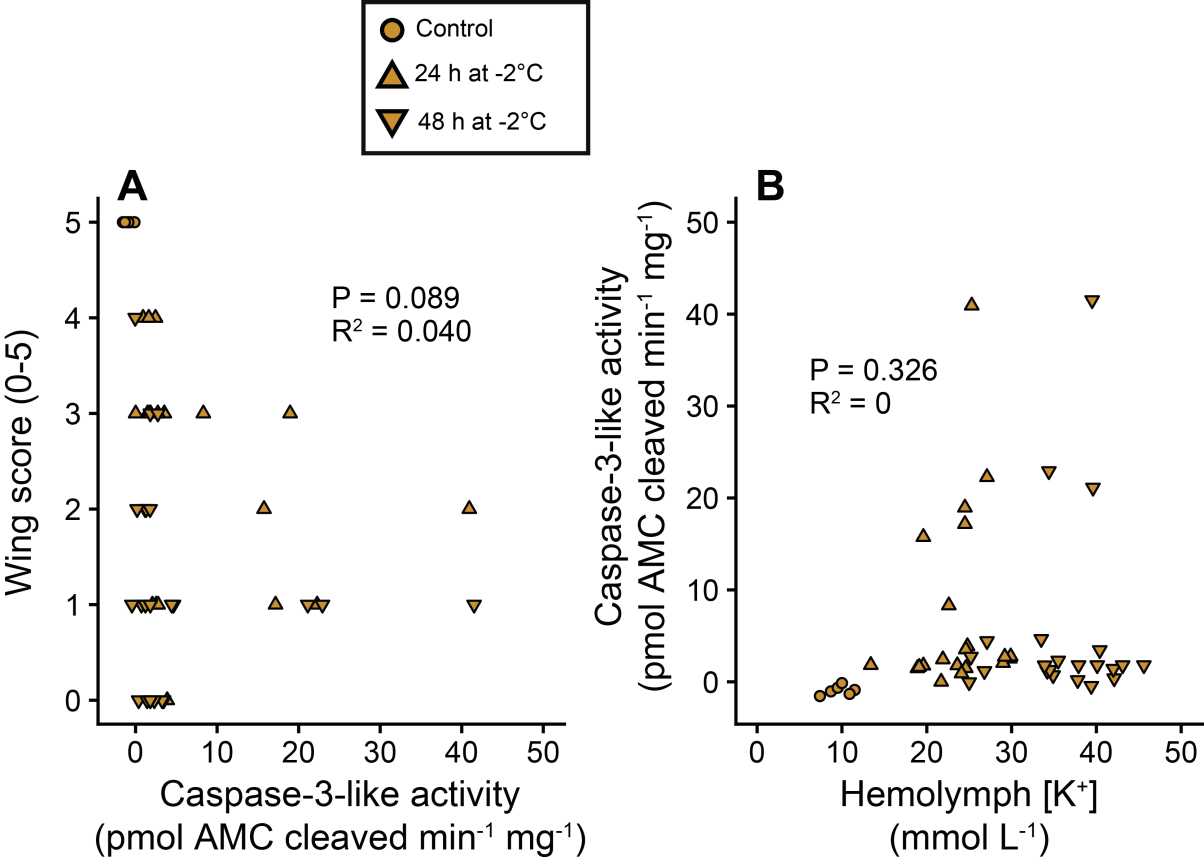

**Fig. S1. Correlations between muscle caspase activity and wing score as well as hemolymph [K^+^] and muscle caspase activity.** Using linear regression we found that (A) the correlation between muscle caspase-3-like activity and the wing score was better than that for organismal survival score (see Fig. 2E) despite still not reaching statistical significance. Similarly, (B) the correlation between hemolymph [K^+^] and muscle caspase-3-like activity was not statistically significant.

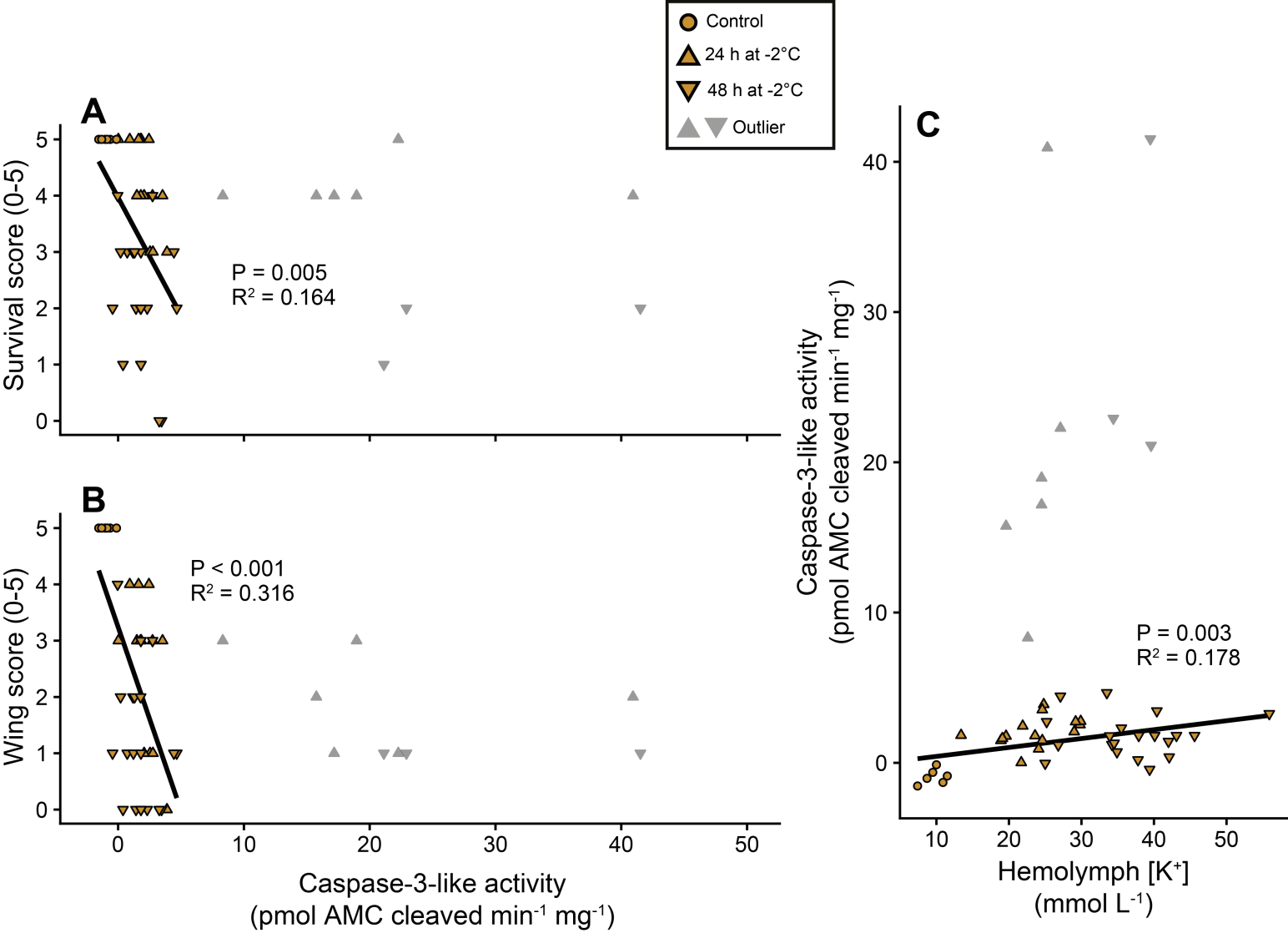

**Fig. S2. Correlations between survival score estimated, muscle caspase-3 activity and hemolymph [K^+^] with outliers removed.** After outliers (grey) was removed using Grubb’s test (which works by sequentially removes outliers) both correlations between (A) survival score, (B) wing score and muscle caspase-3-like activity become statistically significant. Similarly does the correlation between (C) hemolymph [K^+^] and muscle caspase-3-like activity. Despite the statistical significance, the model hold little predictive power compared to the one provided by the hemolymph [K^+^] vs. survival score correlation (see Fig. 2C).
